# Cross-scale Analysis of Temperature Compensation in the Cyanobacterial Circadian Clock System

**DOI:** 10.1101/2021.08.20.457041

**Authors:** Yoshihiko Furuike, Dongyan Ouyang, Taiki Tominaga, Tatsuhito Matsuo, Atsushi Mukaiyama, Yukinobu Kawakita, Satoru Fujiwara, Shuji Akiyama

## Abstract

Clock proteins maintain constant enzymatic activity regardless of temperature, even though thermal fluctuation is accelerated as temperature increases. We investigated temperature influences on the dynamics of KaiC, a temperature-compensated ATPase in the cyanobacterial circadian clock system, using quasielastic neutron scattering. The frequency of picosecond to sub-nanosecond incoherent local motions in KaiC was accelerated very slightly in a temperature-dependent manner. Our mutation studies revealed that internal motions of KaiC include several contributions of opposing temperature sensitivities. To take advantage of this balancing effect, the motional frequency of local dynamics in KaiC needs to exceed ∼0.3 ps^-1^. Some of the mutation sites may be in a pathway through which the motional frequency in the C-terminal domain of KaiC is fed back to the active site of ATPase in its N-terminal domain. The temperature-compensating ability at the dynamics level is likely crucial for circadian clock systems, into which the clock proteins are incorporated, to achieve reaction- or even system-level temperature compensation of the oscillation frequency.

## Introduction

Circadian clocks are endogenous timing systems that rhythmically control various biological processes with an approximately 24-hour period ^1^. This rhythm persists stably even without any external cues, and the period length is kept constant even when ambient temperature changes (temperature compensation). The phase of the clock system can be shifted upon receiving external stimuli such as light and temperature and then synchronized to the phase of the outer world. These unique characteristics enable organisms to optimize their fitness during day/night environmental cycles ^2, 3, 4, 5^.

Temperature compensation is a remarkable characteristic of the circadian clock systems. Q10 values, the factor by which reaction speed or cycle frequency is accelerated by increasing the ambient temperature by 10°C, are mostly in the range of 0.9–1.1 for circadian clock systems, whereas those of most biochemical reactions ^6^ and the Belousov–Zhabotinsky oscillator ^7, 8, 9, 10^ range from 2 to 3. As schematically shown in the middle panel of Fig. 1*A*, circadian clocks exhibiting system-level temperature compensation often comprise unique clock proteins with temperature-compensated biochemical activities such as ATPase ^11, 12, 13^ and kinase/phosphatase ^14, 15, 16^. A simple but attractive idea that emerged from studies ^12, 13, 15, 17, 18^ of these biochemical activities is that the reaction-level temperature compensability is somehow correlated to system- or even higher cell-level temperature compensability (lower panel of Fig. 1*A*). A great deal of effort has been devoted to elucidating the mechanism of this connectivity via experimental ^12, 13, 15, 16, 18^ and modeling ^19, 20, 21, 22, 23^ approaches.

**Figure 1.**
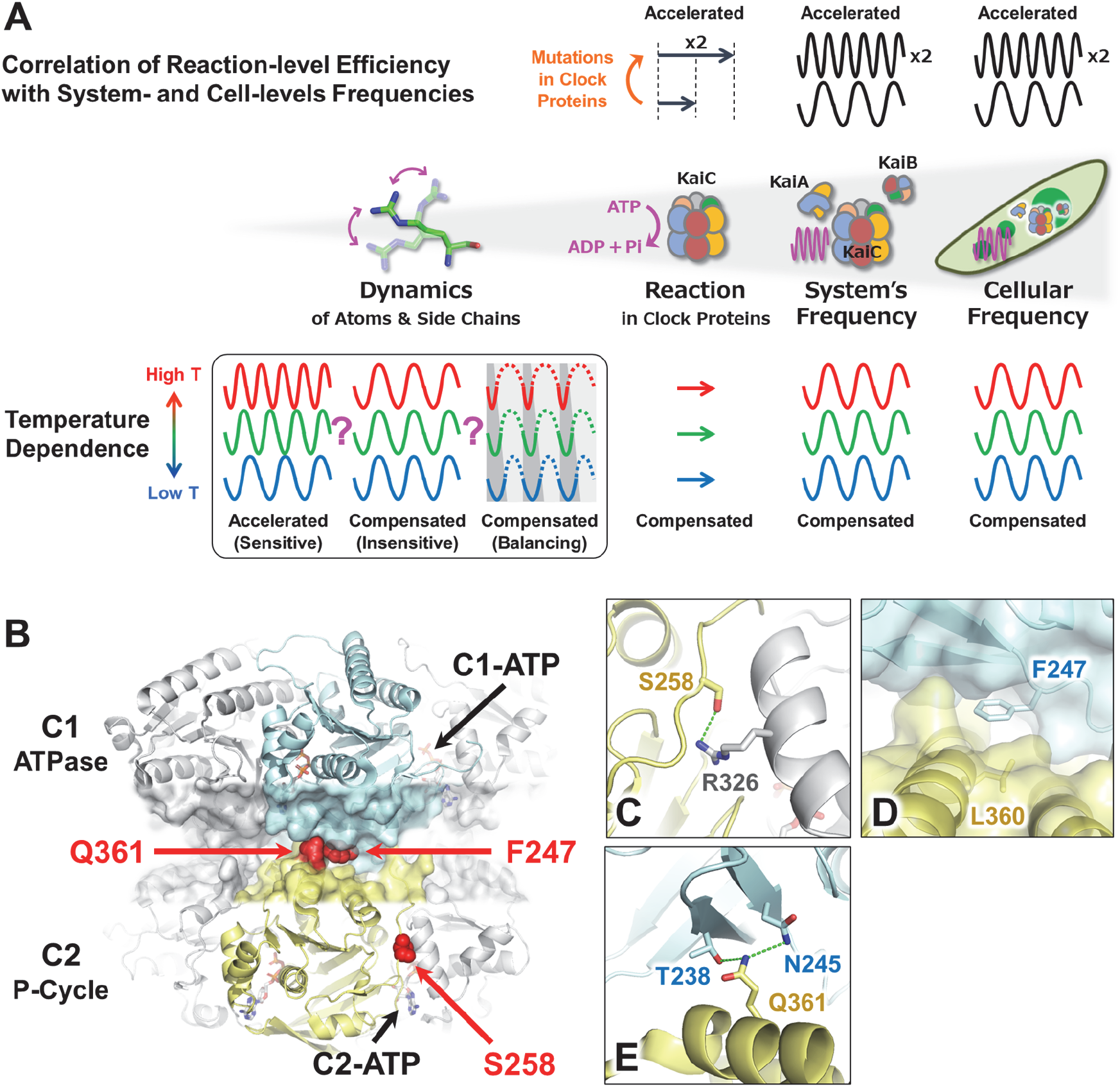
Impact of KaiC dynamics on temperature compensability in the circadian clock system of cyanobacteria. (*A*) Potential cross-scale causal relationship in the circadian clock system of cyanobacteria. (*Middle*) Spatiotemporal hierarchy spanning from atomic-scale dynamics, molecular-scale reaction, molecular system-scale frequency, and cellular-scale frequency. (*Upper*) Correlation of the reaction-level efficiency with system- and cell-level frequencies in cyanobacteria. When the ATPase activity of KaiC doubles as a result of amino acid substitutions (orange arrow), both the *in vitro* system- and cell-level frequencies also double ^12, 17, 25^. (*Lower*) Potential correlation of temperature sensitivity of clock protein dynamics with reaction-, system-, or cell-level temperature compensability. In cyanobacteria, the temperature sensitivity of ATPase is correlated with those of the system and cell levels. (*Box*) Three extreme cases of temperature influence on clock-protein dynamics. Thermal fluctuations are sensitively accelerated (*Left*), insensitively compensated (*Center*), or sensitively compensated through a balance of opposing contributions (*Right*). (*B*) Temperature-dependent ATPase mutation sites mapped onto the crystal structure of KaiC ^32^. Zoomed-in-views of (*C*) S258, (*D*) F247, and (*E*) Q361 in KaiC. Hydrogen bonds are highlighted by green dotted lines.

Nevertheless, temperature compensation remains a puzzling phenomenon in terms of protein dynamics, as atoms and side chains in proteins should fluctuate more frequently at higher temperature due to their greater thermal energy. Three extreme cases can be considered (box in Fig. 1*A*). First, thermal fluctuation in these key clock proteins is accelerated in a temperature-dependent manner, as observed for ordinary proteins. Second, the atoms and side chains in temperature-compensated clock proteins fluctuate in a temperature-insensitive manner via unknown mechanisms. Third, some opposing but balancing contributions act at the level of protein dynamics level, i.e., one contribution accelerates but the other decelerates so that they differentially affect the elementary reaction steps to achieve overall compensation. Experimental investigations of the dynamics of temperature-compensated clock proteins are needed to distinguish among these cases.

KaiC, a temperature-compensated clock protein in cyanobacterium *Synechococcus elongatus* PCC 7942 ^11, 12, 13^, acts as a circadian oscillator with other two clock proteins, KaiA and KaiB ^24^. Its most striking feature is that its temperature-compensated circadian rhythm can be reconstructed even *in vitro* by mixing KaiA, KaiB, and KaiC in the presence of ATP ^25^ (Fig. 1*A*). An ATP molecule binds a Walker motif in each of the tandemly duplicated domains called the N-terminal C1 and C-terminal C2 domains (Fig. 1*B*). These ATP binding events trigger oligomerization of KaiC into a double-ring hexamer. The ATP molecule bound to the C2 domain (C2-ATP) is used mostly as the source of phosphoryl group that is transferred to (auto-kinase) and then removed from (auto-phosphatase) Ser431 and Thr432 in KaiC. In the presence of KaiA and KaiB, the status of the dual phosphorylation site alters in a cyclic manner: ST → SpT → pSpT → pST → ST, where S, T, pS, and pT represent Ser431, Thr432, phosphorylated Ser431, and phosphorylated Thr432, respectively ^26, 27^. The frequency of this phosphorylation cycle (P-cycle) is proportional to the rate of hydrolysis of C1-bound ATP (C1-ATP) in the absence of KaiA and KaiB (upper panel of Fig. 1*A*) ^12, 13, 17^. More importantly, the ATPase activity of KaiC is perfectly temperature-compensated (Q10_ATP_ = 1.0) ^12^. The *in vitro* Kai oscillator that consists of temperature-compensated KaiC provides a practical means for studying cross-scale properties of temperature-compensation phenomena at the system, reaction, and dynamics levels (Fig. 1*A*).

Quasielastic neutron scattering (QENS) is a powerful and direct technique for accessing protein dynamics at the picosecond to nanosecond time scales ^28^. Because hydrogen atoms, which constitute up to half of all atoms in proteins, are distributed near uniformly in the three-dimensional structures of proteins, an averaged view of protein dynamics can be extracted from QENS spectra ^29^. In this study, we designed a series of KaiC mutants (Fig. 1*B*) with temperature-dependent periods (Fig. 1*C*-*E*), and measured the temperature dependence of their protein dynamics using the neutron spectrometer DNA at J-PARC MLF ^30^. Our QENS data revealed that KaiC establishes reaction-to cell-level temperature compensation by keeping the frequency of its picosecond to sub-nanosecond incoherent motions nearly temperature-insensitive, but still high enough to exceed certain threshold limits.

## Results

### Screening and Characterization of Temperature-dependent Mutants of KaiC

As confirmed by the positions along the horizontal axis in Fig. 2*A*, the ATPase activity of KaiC^WT^ is as low as 11 d^-1^ and almost temperature-insensitive (Q10_ATP_^WT^ = 0.89 ± 0.10), as previously reported ^12^. Consistently with previous studies ^11, 15, 25^, the P-cycle frequency (*f*_P_ = 24 / period) of KaiC^WT^ in the presence of KaiA and KaiB was also temperature-compensated (Fig. 2*B*) (Q10_fp_^WT^ = 1.08 ± 0.01). Consequently, data points taken at different temperatures for KaiC^WT^ nearly overlapped in the ATPase – *f*_P_ plot (Fig. 2*A*). Taking advantage of potential reaction–system correlations, we screened the temperature-dependent mutants of KaiC using an *in vitro* ATPase-based screening system ^31^. Among a number of KaiC mutants screened for temperature-dependent ATPase activity, three candidates were characterized in detail as they revealed stable but temperature-dependent system-level oscillation (Fig. 2*C*–*E*).

**Figure 2.**
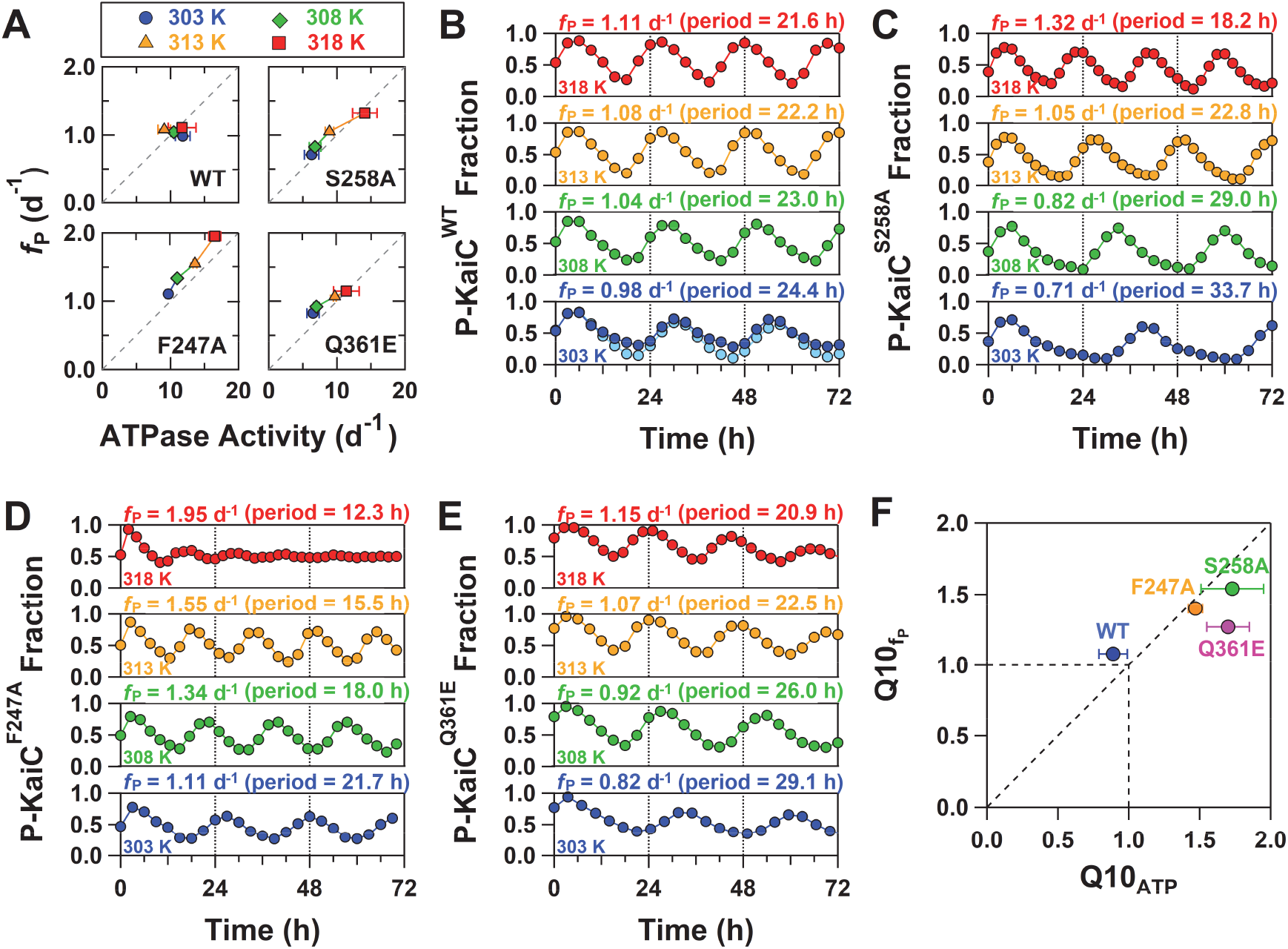
Screening and characterization of temperature-dependent mutants of KaiC. (*A*) Temperature dependence of the relationship between the ATPase activity of KaiC alone and *in vitro* KaiC phosphorylation-cycle (P-cycle) frequency (*f*_P_ = 24 / period) in the presence of KaiA and KaiB. P-cycles of (*B*) KaiC^WT^, (*C*) KaiC^S258A^, (*D*) KaiC^F247A^, and (*E*) KaiC^Q361E^ at four different temperatures: 303 (blue), 308 (green), 313 (orange), and 318 K (red). Light blue circles plotted in panel (*B*) correspond to the P-cycle of KaiC^WT^ in a D_2_O buffer. (*F*) Correlation of Q10 values between ATPase (Q10_ATP_) and P-cycle (Q10_fp_).

The first was the S258A mutant of KaiC (KaiC^S258A^). According to the X-ray crystal structure of KaiC^WT 32^, S258 is located in the C2 domain (Fig. 1*B* and 1*C*). The ATPase activity of KaiC^S258A^ at 303 K (6.3 ± 1.1 d^-1^) was lower than that of KaiC^WT^ (11.8 ± 1.1 d^-1^) but increased in a temperature-dependent manner up to 14.1 ± 1.8 d^-1^ at 318 K (Q10_ATP_^S258A^ = 1.73 ± 0.22) (Fig. 2*A*). Consistent with this, the *f*_P_ value for KaiC^S258A^ increased from 0.71 to 1.32 d^-1^ as the temperature increased (Fig. 2*C*) (Q10_fp_^S258A^ = 1.54 ± 0.01). Because of this correlation, the data trace for KaiC^S258A^ extends almost diagonally from low-to high-temperature conditions in the ATPase – *f*_P_ plot (Fig. 2*A*). The two mutants (KaiC^F247A^, KaiC^Q361E^), which replaced residues that neighbored in the C1–C2 interface (Fig. 1*B*), are also traced diagonally (Fig. 2*A*). The *f*_P_ value for KaiC^F247A^ increased from 1.11 to 1.95 d^-1^ as temperature increased (Fig. 2*D*) (Q10_fp_^F247A^ = 1.40 ± 0.01), as observed for the mutant’s ATPase activity (Fig. 2*A*) (Q10_ATP_^F247A^ = 1.47 ± 0.05), whereas KaiC^Q361E^ exhibited a slightly weakened correlation between Q10_ATP_^Q361E^ (1.70 ± 0.15) and Q10_fp_^Q361E^ (1.27 ± 0.02) (Fig. 2*E* and 2*F*).

Taking into consideration the limited persistence of the KaiC^F247A^ P-cycle at 318 K (Fig. 2*D*), we dissolved KaiC^WT^ and the three KaiC mutants exhibiting reaction–system correlation in a D_2_O buffer and subjected them to QENS experiments at low and high temperatures: 302 and 312 K for the fully dephosphorylated form of KaiC^WT^ (*SI Appendix*, Fig. S1), and 302 and 309 K for KaiC^S258A^, KaiC^F247A^, and KaiC^Q361E^. Noted that the circadian rhythm of KaiC^WT^ was barely influenced by the D_2_O buffer used in this study (light blue circles in Fig. 2*B*).

### KaiC Dynamics Detected by QENS

To investigate the temperature dependence of KaiC dynamics, we conducted QENS experiments at low and high temperatures. As shown in Fig. 3*A*, a difference QENS spectrum *S*(*Q,E*) of KaiC^WT^, where *Q* is the momentum transfer and *E* is the energy transfer of neutrons, could be obtained at each temperature by subtracting the background spectrum of the D_2_O solvent from the sample spectrum, on the basis of scaling factors calculated from neutron scattering cross sections. The resultant *S*(*Q,E*) for KaiC^WT^ at 302 and 312 K shared a common feature of a broadened elastic peak and a quasielastic component derived from global and local motions, respectively (Fig. 3*A*). To analyze the *Q*- and temperature dependencies of these two components quantitatively, we attempted to fit the following equation ^28^ to each *S*(*Q,E*):

**Figure 3.**
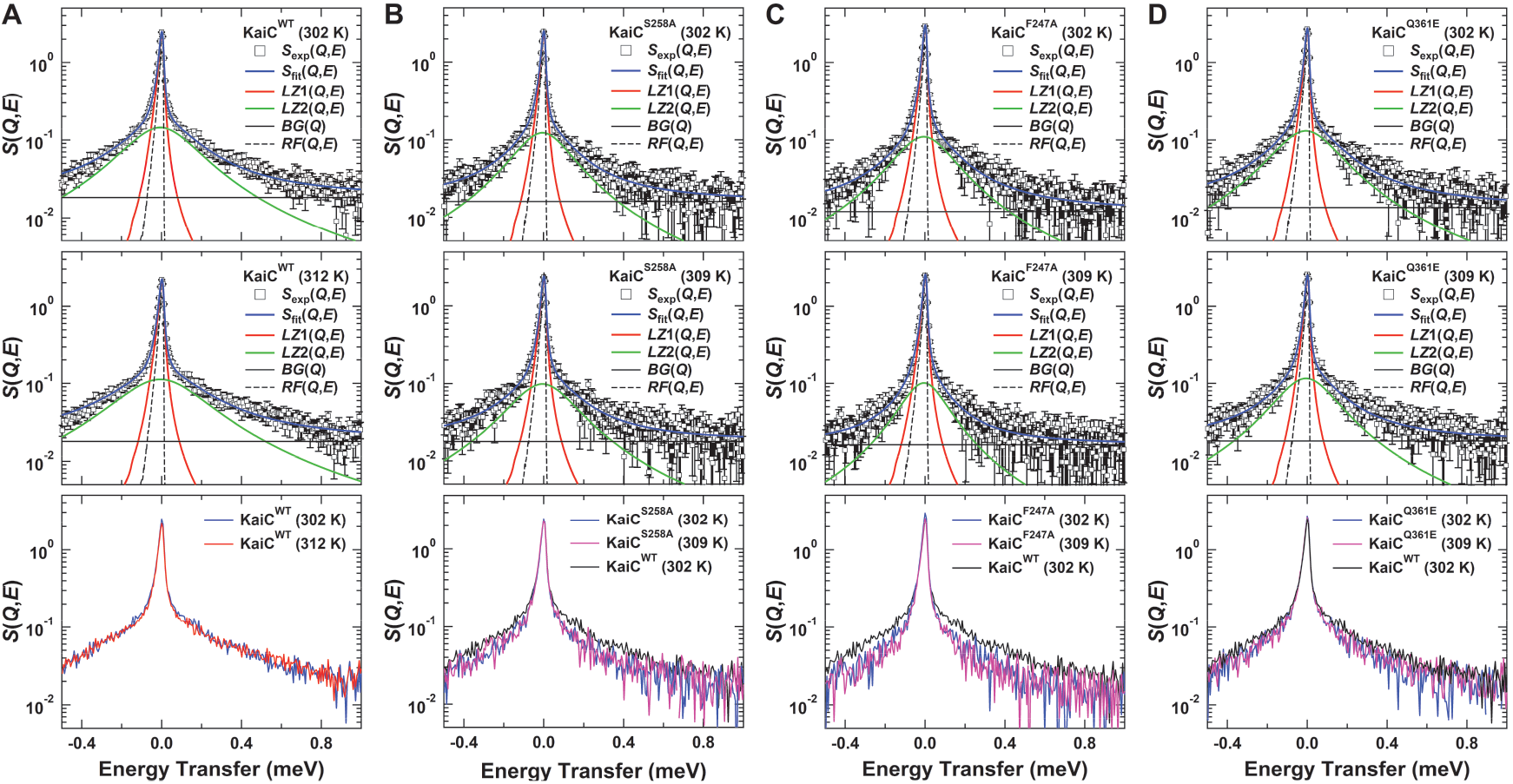
Representative QENS spectra, *S*(*Q,E*), of (*A*) KaiC^WT^, (*B*) KaiC^S258A^, (*C*) KaiC^F247A^, and (*D*) KaiC^Q361E^ at *Q* = 1.45 Å^-1^. (*Upper*) At the low temperature of 302 K. (*Middle*) At a high temperature of 312 or 309 K. Experimental spectra, *S*_exp_(*Q,E*), are fitted using *S*_fit_(*Q,E*), which includes contributions of two Lorentzian functions, *LZ*1(*Q,E*) = *L*_global_(*Q,E*) and *LZ*2(*Q,E*) = *L*_global_(*Q,E*)⊗*L*_local_(*Q,E*), background *BG*(*Q*), and resolution function *RF*(*Q,E*) as defined in Eq. **1**. (*Lower*) Comparison of QENS spectra acquired at low and high temperatures. Blue, red, and magenta lines correspond to the spectra at 302, 312, and 309 K, respectively.

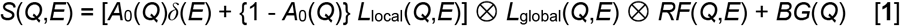

where *A*_0_(*Q*) is the elastic incoherent structure factor (EISF); *δ*(*E*) is Dirac delta function; *RF*(*Q,E*) is the instrumental resolution function obtained from the spectrum of vanadium; *BG*(*Q*) is the background; and *L*_local_(*Q,E*) and *L*_global_(*Q,E*) are Lorentzian functions, (Γ(*Q*) / π) (*E*^2^ + Γ(*Q*)^2^)^-1^, describing local internal motion and global diffusive motion of the protein, respectively, where Γ(*Q*) is the half width at half maximum (HWHM). *Q*-averaged *χ*^2^ values resulting from fitting of Eq. **1** to the experimental data of KaiC^WT^ was ∼ 0.96 ± 0.04 (upper and middle panels of Fig. 3*A*), assuring optimum quality of the curve-fitting procedure. In the following, we describe the global and local motions of KaiC on the basis of temperature dependencies of Γ_global_(*Q*) and Γ_local_(*Q*), respectively.

### Global Motions of KaiC

Γ_global_(*Q*) provides information on the frequency of the global motions, whose properties are most straightforwardly inspected using the plot of Γ_global_(*Q*) vs *Q*^2^. Linear *Q*^2^-dependencies of Γ_global_(*Q*) were confirmed for KaiC^WT^ at both high and low temperatures (Fig. 4*A*), indicating its boundary-free diffusions including translational and rotational diffusions. Thus, the linear slope in Fig. 4*A* relates to an apparent boundary-free diffusion coefficient (*D*_global_). The *D*_global_ for KaiC^WT^ at 302 and 312 K were 3.53 ± 0.06 and 4.32 ± 0.06 10^−7^ cm^2^ s^-1^, respectively (Fig. 4*E*), which were similar to those (3.81 and 4.88 10^−7^ cm^2^ s^-1^) calculated using the known crystal structure of the KaiC hexamer (Fig. 1*B*) (*SI Appendix*, Table S1 and Section S.1.). The observed temperature dependencies of the *D*_global_ (Q10_global_) for KaiC^WT^ and the KaiC mutants (Fig. 4*B*-*D*) were approximately 1.2 (Fig. 4*E*), similar to other examples: ∼1.2 (300–310 K) in human hemoglobin (Hb) ^33^ and ∼1.2 (290–300 K) in α-synuclein (αSyn) ^34^. Thus, KaiC^WT^ and the KaiC mutants were maintained as intact hexamers during the QENS measurements, and underwent the same translational and rotational diffusions as ordinary proteins.

**Figure 4.**
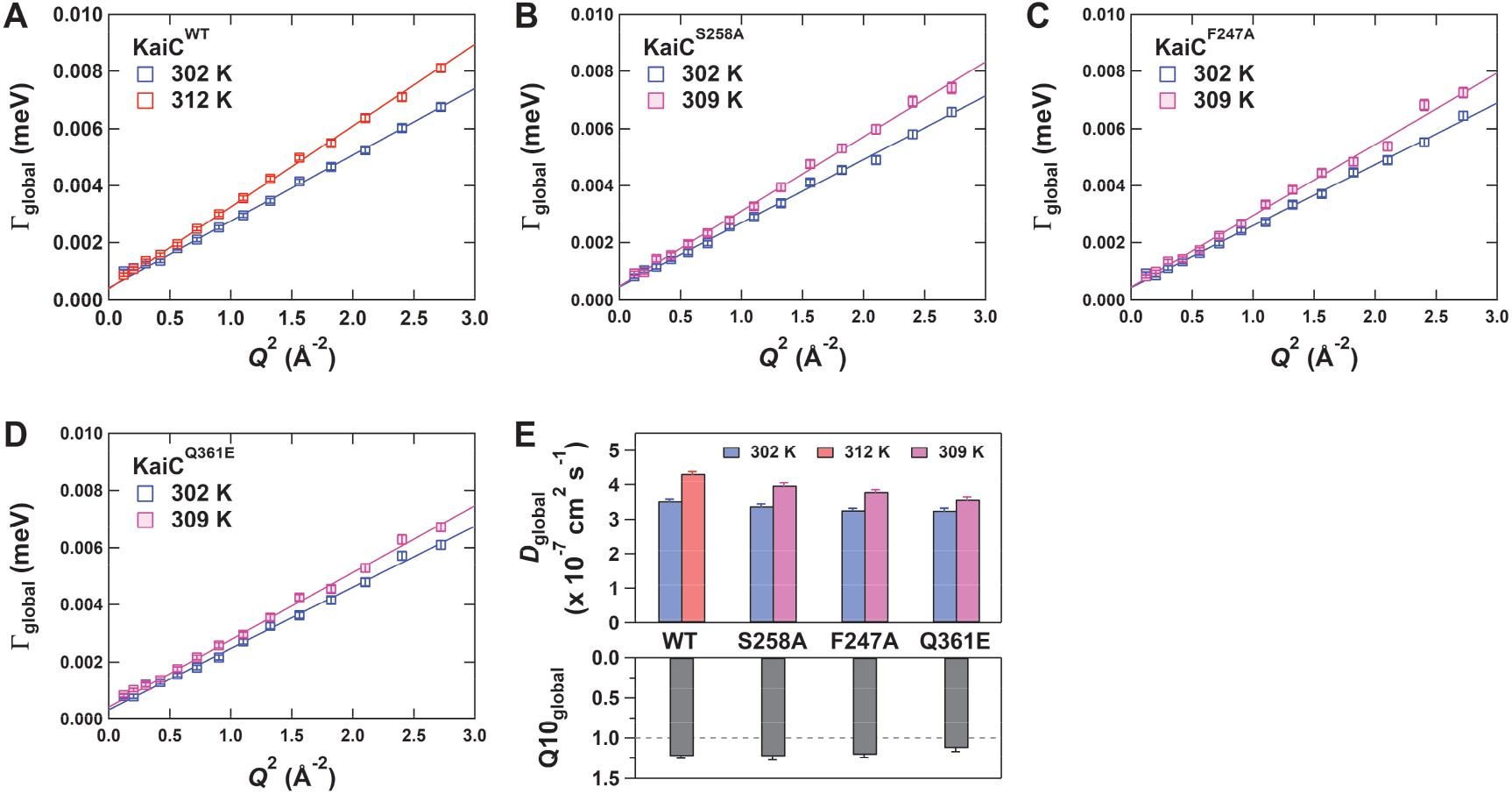
Temperature dependence of global motions in KaiC. *Q*^2^-dependence of Γ_global_(*Q*) for (*A*) KaiC^WT^, (*B*) KaiC^S258A^, (*C*) KaiC^F247A^, and (*D*) KaiC^Q361E^. Blue, red, and magenta boxes correspond to data acquired at 302, 312, and 309 K, respectively. The slope of each linear fit corresponds to the apparent diffusion coefficient, *D*_global_. (*E*) *D*_global_ and its temperature dependence, Q10_global_.

### Balanced Contributions of Local Motions Revealing Different Sensitivities to Temperature

In contrast to the global motions, the local fluctuations of side-chain and main-chain in KaiC revealed unique temperature dependencies. Γ_local_(*Q*) of KaiC^WT^ increased asymptotically to approach a plateau at high *Q*^2^ (Fig. 5*A*). As at high *Q*, the motions over short distances predominate, the plateau values correspond to the elementary displacements of the local motions. Thus, in the saturated *Q*^2^-range higher than 1.8 Å^-2^, a *Q*-averaged ratio of Γ_local_(*Q*) taken at high and low temperatures serves as a good model-free measure of the temperature dependence of the internal motions (Q10_local,app_). The Q10_local,app_ value for KaiC^WT^ was 1.15 ± 0.03 (Fig. 5*E*), implying a limited temperature dependence of the local motions. On the other hand, the temperature-dependent mutants exhibited diverse behaviors in response to temperature change. Near-perfect insensitivities of Γ_local_(*Q*) were observed for both KaiC^S258A^ (Fig. 5*B*, Q10 _local,app_^S258A^ = 1.06 ± 0.07) and KaiC^Q361E^ (Fig. 5D, Q10_local,app_^Q361E^ = 0.96 ± 0.05). Γ_local_(*Q*) of KaiC^F247A^ decreased in an unusual manner as the temperature increased (Fig. 5*C*), indicating an inverse temperature dependence (Q10_local,app_^F247A^ = 0.80 ± 0.06) (Fig. 5*E*). Noted that these unique responses of the KaiC mutants could be visually confirmed by some difference in the raw (lower panel in Fig. 3) and *Q*-averaged (*SI Appendix*, Fig. S2) QENS spectra.

**Figure 5.**
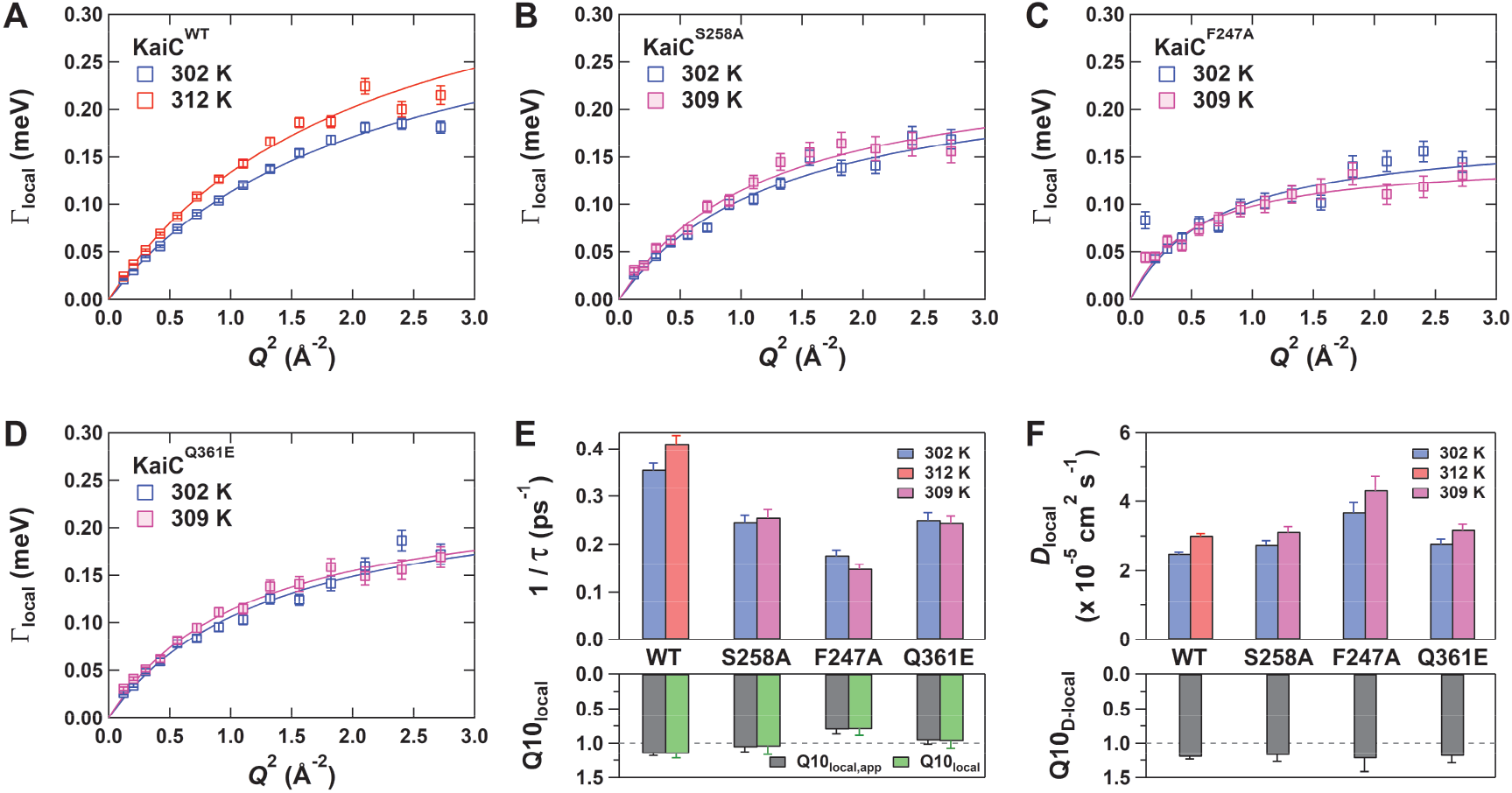
Temperature dependence of local motions in KaiC. *Q*^2^-dependence of Γ_local_(*Q*) for (*A*) KaiC^WT^, (*B*) KaiC^S258A^, (*C*) KaiC^F247A^, and (*D*) KaiC^Q361E^. Blue, red, and magenta boxes correspond to the data acquired at 302, 312, and 309 K, respectively. Solid lines represent resultant fits using a jump-diffusion model that predicts the jump-diffusion coefficient, *D*_local_, as the curvature of the saturating curves and the reciprocal of residence time, *τ*^-1^, as Γ_local_(*Q*) converged at infinite *Q*. (*E*) *τ*^-1^ and its temperature dependence, Q10_local_ (green bars). Q10_local,app_ (gray bars) represents a *Q*-averaged ratio of Γ_local_(*Q*) taken at high and low temperatures using a *Q*^2^-range higher than 1.8 Å^-2^. (*F*) *D*_local_ and its temperature dependence, Q10_D-local_.

The uniqueness of the local motions was also predicted by model-based analysis using a jump-diffusion model ^28^: Γ_local_(*Q*) = *D*_local_*Q*^2^ (1 + *D*_local_*Q*^2^*τ*)^-1^, where *D*_local_ is the jump-diffusion coefficient and *τ* is the residence time spent on one site before jumping to others. The model gave reasonable fits to temperature- and *Q*^2^-dependent Γ_local_(*Q*) (lines in Fig. 5*A*–*D*). The resultant *τ*^-1^ values for KaiC^WT^ at 302 and 312 K were 0.36 ± 0.01 and 0.41 ± 0.02 ps^-1^, respectively (Fig. 5*E*). The model-based temperature dependence (Q10_local_) estimated as a ratio of the *τ*^-1^ values, was 1.15 ± 0.06 for KaiC^WT^, consistent with model-free Q10_local,app_^WT^ of 1.15 ± 0.03 (Fig. 5*E*). The coincidences between Q10_local_ and Q10_local,app_ were also confirmed for the KaiC mutants.

A common feature of the temperature-dependent mutants was substantial deceleration of local dynamics at both temperatures (Fig. 5*E*). The *τ*^-1^ values for KaiC^S258A^ were 0.25 ± 0.01 and 0.26 ± 0.02 ps^-1^ at low and high temperatures, respectively, and were reduced approximately 60– 70% relative to KaiC^WT^ (Fig. 5*E*). Similarly, the local motions in KaiC^Q361E^ was 30∼40% slower (0.24∼0.25 ± 0.01 ps^-1^) than KaiC^WT^ (Fig. 5*E*). Effects of the F247A substitution on local dynamics became so obvious at high temperature as to reveal a reduced *τ*^-1^ value of 0.15 ± 0.01 ps^-1^ (Q10_local_^F247A^ = 0.80 ± 0.09). To ensure temperature compensability at both the reaction and system levels (Fig. 1*A*), the local motions of KaiC must be accelerated very slightly in a temperature-dependent manner (Q10_local_^WT^ = 1.15 ± 0.06), but the absolute frequency of the fluctuations must also exceed certain threshold limits (∼0.3 ps^-1^ in Fig. 5*E*). *D*_local_ was essentially unaffected by temperature change or mutations (Fig. 5*F*), except in the case of KaiC^F247A^, where a large error of *D*_local_ at high temperature prevented us from identifying clear tendency.

These results suggest that KaiC dynamics include temperature-dependent acceleration (Q10_local_^WT^ = 1.15 ± 0.06) as well as temperature-independent (Q10_local_^S258A^ = 1.05 ± 0.11, Q10_local_^Q361E^ = 0.97 ± 0.10) and temperature-dependent decelerating motions (Q10_local_^F247A^ = 0.80 ± 0.09) (Fig. 5*E*). The fact that even a single amino acid substitution has significant impact on motional frequency and its temperature sensitivity (Fig. 5*E*), depending on where it is introduced, implies that overall temperature compensability of the local motions in KaiC is in a delicate balance that relies on the interplay of several opposing temperature sensitivities.

### Fractional Increase in Apparently Immobile Atoms in the Temperature-dependent ATPase Mutants of KaiC

EISF is defined as the ratio of the elastic peak intensity to the sum of elastic and quasielastic scattering intensities as in Eq. (1), providing information on the geometry of molecular motions and the fraction of mobile and immobile atoms on the time scale (∼55 ps) of the spectrometer. As shown in Fig. 6*A*–*D*, EISF of KaiC^WT^ and the mutants were plotted against *Q* and then fitted using the following equation, assuming diffusion in an ensemble of spheres whose radii (*a*) follow a lognormal distribution ^35^:

**Figure 6.**
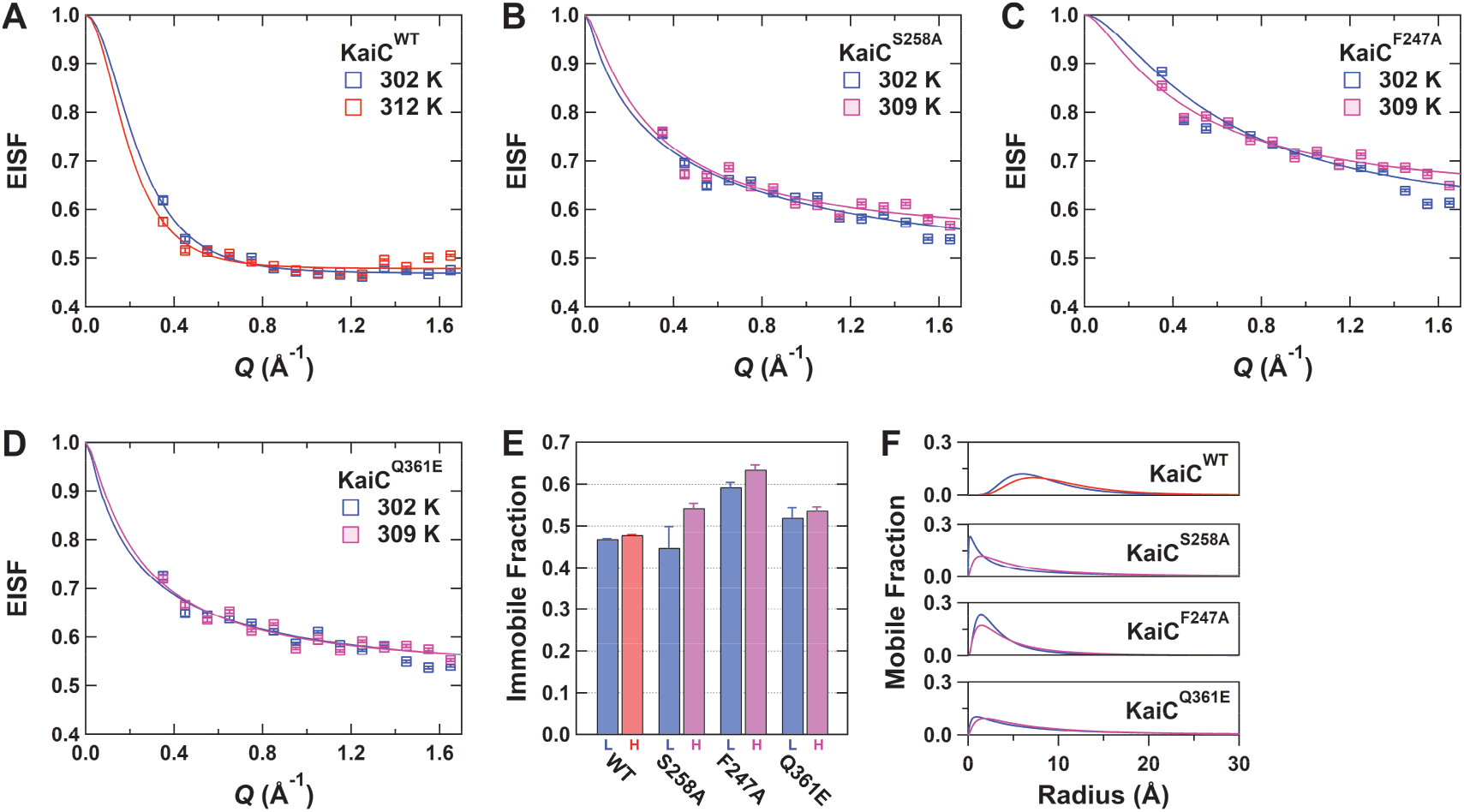
EISF analyses under the assumption of sphere-ensemble model for (*A*) KaiC^WT^, (*B*) KaiC^S258A^, (*C*) KaiC^F247A^, and (*D*) KaiC^Q361E^. Blue, red, and magenta boxes correspond to data acquired at 302, 312, and 309 K, respectively. Solid lines represent the resultant fits of Eq. **2**. (*E*) Immobile fraction, *p*. Blue- and red/magenta-colored bars correspond to the parameters for low (L) and high (H) temperatures, respectively. (*F*) Radial distribution functions of mobile fraction at low (blue) and high (red or magenta) temperatures.

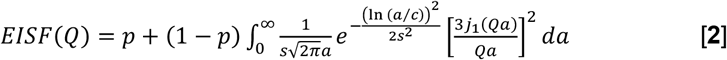

where *p* is the fraction of atoms whose motions are outside the current instrumental energy window and therefore appear immobile; (1 - *p*) correspond to the fraction of atoms diffusing within the sphere ensemble; *c* is the median of the distribution; *s* is the variance in the natural logarithmic space; and *j*_1_ denotes the spherical Bessel function of the first kind of order.

The results of EISF analysis support the unique behaviors of the temperature-dependent mutants of KaiC. Consistent with the Q10_local_ value of 1.15 ± 0.06 for KaiC^WT^, its immobile fraction was maintained at approximately 0.47 at both high and low temperatures (Fig. 6*E*). The immobile fractions of KaiC^S258A^ and KaiC^Q361E^ were nearly unaffected by temperature change within the experimental error. On the other hand, the immobile fraction of KaiC^F247A^, whose local motions exhibited an inverse temperature dependence (Q10_local_^F247A^ = 0.80 ± 0.09), increased 7% as the temperature increased. At the same time, all the KaiC mutants exhibited larger immobile fractions than KaiC^WT^. This result that the amino acid substitutions shifted certain motions from mobile to immobile fractions (Fig. 6*E*) is consistent with the systematic decrease in the *τ*^-1^ values in the KaiC mutants (Fig. 5*E*). A similar trend was also confirmed in the EISF analysis based on a diffusion-inside-two-spheres model ^36^ (*SI Appendix*, Fig. S3 and Section S.2.). In KaiC^WT^, atoms diffusing within a radius of 6–7 Å (Fig. 6*F*) constituted a major mobile component at low and high temperatures. By contrast, the mobile fractions of the KaiC mutants were distributed mainly in reduced radii of 2–3 Å relative to KaiC^WT^. Thus, KaiC^S258A^, KaiC^F247A^, and KaiC^Q361E^ are the temperature-dependent ATPase mutants with reduced frequencies (Fig. 5*E* and 6*E*) and amplitudes (Fig. 6*F*) of local motions, and likely behave as if they sense temperatures lower than that felt by KaiC^WT^.

## Discussion

Over the past decades, chronobiologists have sought a reasonable model that explains the three physiological properties of the circadian clock systems: self-sustained oscillation, temperature compensation, and synchronization ^2, 3, 4^. The circadian clock of cyanobacteria is an ideal experimental system for this purpose, as its physiological properties can be studied in relation to the physicochemical properties of clock-related components at the molecular and atomic scales.

Among the three Kai proteins, KaiC is the core of the cyanobacterial clock system. In the presence of both KaiA and KaiB, KaiC exhibits a phosphorylation rhythm (Fig. 2*B*) whose frequency (*f*_P_) is correlated with the ATPase activity of KaiC alone (Fig. 2*A*). For example, when the ATPase activity of KaiC doubles as a result of amino acid substitutions, the frequencies of both the *in vitro* system-scale and the cellular-scale rhythms also double (upper panel of Fig. 1*A*) ^12, 17, 25^.

This causal relationship, in which properties are transferred through upward causation from bottom to top in the spatiotemporal hierarchy, was also confirmed for temperature compensation from the reaction to the cellular scale (lower panel of Fig. 1*A*). The ATPase activity of KaiC^WT^ is temperature compensated; probably because of this, both the *in vitro* and *in vivo* rhythms ^13^ are also temperature-independent. This phenomenological interpretation is further supported by the three examples of temperature-dependent mutants of KaiC. In contrast to KaiC^WT^ (Q10_ATP_^WT^ = 0.89 ± 0.10), the *f*_P_ values for KaiC^S258A^ (Q10_ATP_^S258A^ = 1.73 ± 0.22), KaiC^F247A^ (Q10_ATP_^F247A^ = 1.47 ± 0.05), and KaiC^Q361E^ (Q10_ATP_^Q361E^ = 1.70 ± 0.15) increased in a temperature-dependent manner (Fig. 2*A*). These results clearly demonstrate that temperature compensation is established from the reaction to the system scale through cross-scale causal relationships arising from the temperature-compensated ATPase activity of KaiC.

Our QENS observations indicate that the causal relationship becomes less simple when the spatiotemporal scale is extended down to local dynamics (*Box* in Fig. 1*A*). This is because the local motions in KaiC^WT^ and its temperature-dependent ATPase mutants respond uniquely to temperature increase. Taking into account the fact that temperature increase slightly accelerates local dynamics of KaiC^WT^ (Q10_local_^WT^ = 1.15 ± 0.06), the crosstalk between the reaction and dynamics levels is not so simple, even if the causal relationship can be extended to down to the microscopic scale. In fact, the simple extension is unrealistic for KaiC^F247A^, as its local fluctuation obeys an inverse temperature dependence (Q10_local_^F247A^ = 0.80 ± 0.09) as opposed to its temperature-dependent ATPase activity (Q10_ATP_^F247A^ = 1.47 ± 0.05). The same applies to both KaiC^S258A^ and KaiC^Q361E^, whose local dynamics are temperature-compensated (Q10_local_^S258A^ = 1.05 ± 0.11, Q10_local_^Q361E^ = 0.97 ± 0.10).

These puzzling yet important observations may be reasonably interpreted by taking into account the absolute frequencies of the local fluctuations. Apart from the temperature sensitivity, all the temperature-dependent mutants of KaiC exhibited a substantial deceleration of local dynamics at both temperatures (Fig. 5*E*). Our observations suggest that KaiC^WT^ establishes temperature compensation at both the reaction and system levels by keeping the frequencies of its local motions nearly temperature-insensitive but high enough to exceed certain threshold limits (∼0.3 ps^-1^). A rate-limiting step of the ATPase cycle, which is yet to be determined experimentally, likely receives some feedback through local protein motions to achieve a constant ATPase activity; therefore, the frequency of the associated fluctuations must be similar regardless of temperature (Fig. 7*A*). The QENS data of both KaiC^S258A^ and KaiC^Q361E^ imply that even if motional frequency is nearly constant when temperature changes, the feedback frequency through the local motions must be high enough to suppress the inherent temperature-dependent increase (Fig. 5*E*).

**Figure 7.**
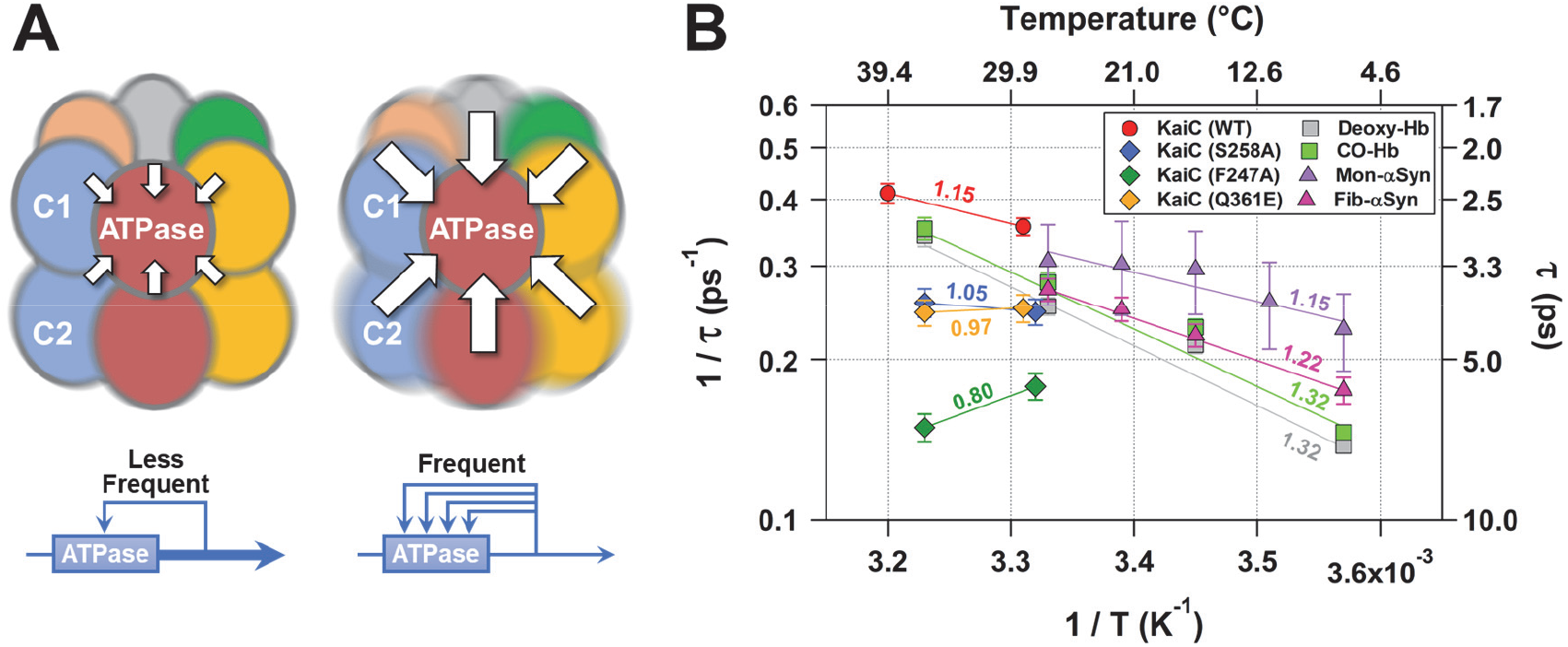
Temperature-insensitive but active internal motions in KaiC as the basis of temperature compensability in the circadian clock system of cyanobacteria. (*A*) A model for C1-ATPase receiving regulatory feedback through thermal fluctuations of KaiC. (*Left*) Temperature-dependent ATPase under less frequent feedback. (*Right*) Temperature-compensated ATPase under frequent feedback. (*B*) Temperature dependences of jump-diffusion frequency (*τ*^-1^) for KaiC, human hemoglobin (Hb), and α-synuclein (αSyn). QENS data of Hb and αSyn are taken from previous studies ^33, 51^; activation energies of deoxygenated Hb (deoxy-Hb), CO-bound Hb (CO-Hb), fibrillized αSyn (fib-αSyn), and monomeric αSyn (mon-αSyn) are 5.2 ± 0.3, 5.2 ± 0.3, 3.7 ± 0.7, and 2.6 ± 1.9 kcal mol^-1^, respectively. Values given near fitting lines represent the Q10_local_ values estimated for 303 K.

Our results are worth discussing from the viewpoint of the crystallographic structure. S258 is located in the outer radius side of the C2 ring (Fig. 1*B*); the side chain is hydrogen bonded to that of R326 in the neighboring C2 domain (Fig. 1*C*). This inter-domain hydrogen bond is thus disrupted in KaiC^S258A^. We speculate that inter–C2-domain hydrogen bonds play important roles in keeping the motional frequency of KaiC intact.

Although F247 and Q361 neighbor each other in the C1–C2 interface (Fig. 1*B*), the local dynamics resulting from each mutation obeyed different temperature dependencies (Fig. 5*E*). The fact that mutations into the C1–C2 interface produced variants with diverse temperature dependence may indicate that the C1–C2 interface is inherently dynamic and fluctuating, and is formed by complex interactions that themselves have diverse dependence on temperature. Actually, the C1–C2 interface is stabilized by a limited number of inter-domain contacts with lower packing density than the C1–C1 and C2–C2 interfaces (*SI Appendix*, Fig. S4), exemplified by the non-bonded interactions associated with F247 and Q361 (Fig. 1*B*). The plane of the phenyl ring of F247 in the C1 domain lies nearly parallel to the C1–C2 interface (Fig. 1*D*), filling the loosely packed boundary through a hydrophobic interaction with L360 in the C2 domain. The F247A substitution is thus interpreted as the mutation that converts the local dynamics from nearly temperature-insensitive to inverse temperature-dependent (Q10_local_^F247A^ = 0.80 ± 0.09) by making the intrinsically low-density interface even looser and more fluctuating. This suggests that the lower but delicate packing density at the C1–C2 interface is critical for the temperature-compensated local dynamics in KaiC. Consistent with this explanation, the temperature-compensatory nature of the local motions is maintained for KaiC^Q361E^ (Q10_local_^Q361E^ = 0.97 ± 0.10), whose packing density at the C1–C2 interface is not greatly affected by the replacement of glutamine to glutamate (Fig. 1*E*). However, because the neutral side chain of Q361 in the C2 domain is potentially hydrogen-bonded with either T238 or N245 in the C1 domain (Fig. 1*E*), the Q361E substitution should modulate or even disrupt the C1–C2 hydrogen-bond interactions with minimal impacts on packing density. Thus, the reduced motional frequency in KaiC^Q361E^ can be ascribed to a weakened coupling between the C1 and C2 domains through their local motions. In addition to our findings in this study, several lines of evidence have revealed the dynamic properties of inter-ring stacking of KaiC during the circadian cycle ^37^. In KaiC^WT^, both F247 and Q361 may be positioned in a pathway or a delicate node that transmits the motional frequency of the C2 domain and its temperature dependence to the C1 domain, where the ATPase active site is located (Fig. 1*B* and 7*A*).

Several studies have suggested that fluctuations and structural polymorphs of the clock proteins play important roles in the circadian clock systems. In mammalian systems, CKIδ-dependent phosphorylation is one of the key reactions that regulates period length and its temperature sensitivity ^16^. In CKIδ, the temperature dependence of substrate affinity is compensated by the opposing temperature dependence of product affinity ^18^. On the basis of molecular dynamics (MD) simulation, the authors of that study proposed that the temperature dependence of the amplitude of particular local fluctuations is reversed in the substrate- and product-bound forms, and that this reversal is one of the origins of the biochemical opposition. Structural polymorphs of the clock proteins, which have been confirmed by crystallography, NMR, and MD simulation, also play important roles in the substrate selectivity of CKIδ ^38^ and the interaction between CRY1/2 and the CLOCK-BMAL1 complex ^39^. In the cyanobacterial system, the rhythmic stacking/unstacking of C1- and C2-rings is coupled to the P-cycle of KaiC ^37^. The MD simulation of KaiC suggests a large-scale conformational change during ADP release from the C2 domain ^40^. These previous studies mostly provide information on the amplitudes of internal motions, such as the magnitude and polymorphism of the structural changes. By contrast, this study provided further insights into the frequency of the internal motions and its temperature dependence through a more direct experimental approach. The observation that the internal motions of KaiC consist of temperature-dependent accelerating components as well as temperature-independent and temperature-dependent decelerating components may be reminiscent of the opposing fluctuation amplitudes suggested for CKIδ ^18^. It is important to note, however, that the frequency of the internal motions needs to be high enough to take advantage of the intrinsic balancing effect implemented in KaiC.

To date, the QENS method has been used in many studies to characterize the overall picture of protein dynamics ^29^. The scope of QENS research is expanding from small single-domain proteins ^41, 42^ to larger and more complex molecular systems ^43, 44, 45^, as well as to large-scale conformational changes such as those that occur upon ligand binding ^33, 46, 47^, pressurization ^48, 49, 50^, unfolding ^35, 51, 52, 53, 54, 55^, and fibrillization ^34, 56^.

It is worth discussing our observations in the light of previous QENS studies on proteins other than clock-related proteins. The results reported to date for various protein samples indicate that the *τ* values are mostly distributed in the range of 1–20 ps (*τ*^-1^ = 0.05–1.0 ps^-1^) (see Grimaldo *et al* ^29^ and references therein), although attention must be paid to the differences in measurement temperature, energy resolution, and analysis methods. The *τ* value for KaiC^WT^, 2.4–2.8 ps (*τ*^-1^ = 0.36–0.41 ps^-1^), is included in the above range, indicating that the internal motion of KaiC^WT^ is neither exceptionally slow nor too fast.

More detailed comparisons of internal motions are possible by making reference for QENS data for Hb ^33^ and αSyn ^51^, which were acquired at the same beamline with the same energy resolution (12 μeV / 55 ps). As shown in Fig. 7*B*, the jump-diffusion frequency of KaiC^WT^ at 302 K (*τ*^-1^ = 0.36 ± 0.01 ps^-1^) is the fastest among the four examples (*τ*^-1^ = 0.25–0.31 ps^-1^) for deoxygenated Hb (deoxy-Hb), CO-bound Hb (CO-Hb), a fibril state of αSyn (fib-αSyn), and a monomeric but intrinsically disordered form of αSyn (mon-αSyn). The *τ*^-1^ value for KaiC^WT^ is less temperature-dependent (Q10_local_^WT^ = 1.15 ± 0.06) than those of deoxy-Hb (Q10_local_^deoxy-Hb^ = 1.32 ± 0.02), CO-Hb (Q10_local_^CO-Hb^ = 1.32 ± 0.02), and fib-αSyn (Q10_local_^fib-αSyn^ = 1.22 ± 0.05). At the same time, it is interesting to note that mon-αSyn exhibits a temperature dependence (Q10_local_^mon-αSyn^ = 1.15 ± 0.12) comparable to that of KaiC^WT^. While this similarity of the Q10_local_ value cannot be explained simply in terms of structural similarity, it implies that the C1–C2 interface has a more dynamic nature than would be expected from the static crystal structure (Fig. 1*B* and *SI Appendix*, Fig. S4). In contrast to the insensitivity of the internal dynamics upon an R(CO)-to-T(deoxy) allosteric transition of Hb (Fig. 7*B*), the temperature-dependent mutants of KaiC exhibit dramatic changes in the motional frequency and temperature dependence. Based on these comparisons, we speculate that introducing mutations that disrupt inter-domain interactions may cause some changes in the local motions of Hb.

The mechanism by which a single amino acid substitution can exert a notable effect on internal protein motions deserves further investigation. Given that QENS detects the average image of the fluctuations of all hydrogen atoms scattered throughout the protein molecule, it is rare that a mutation of just one amino acid will be detected as a large change in the QENS spectrum, as observed in this study. This may be related to the fact that KaiC, as the core of the clock system, is required to maintain low enzyme activity independent of temperature. Our observations indicate that KaiC utilizes a subset of internal motions directly or indirectly to control its ATPase activity in the C1 domain so that it does not become more active as temperature increases (Fig. 7*A*). At the same time, there is growing experimental evidence that ordinary enzymes actively utilize internal motions, thereby increasing the efficiency of overall catalytic reactions ^57, 58, 59^. Because functionally relevant fluctuations often refer to collective motions of atoms on slower time scales (μs to ms), care must be taken in discussing those motions in relation to the incoherent and fast (ps) dynamics detected in this study. However, several studies have demonstrated a linkage between ps–ns fluctuations and slower motions associated with catalytic reactions ^60, 61, 62^. Therefore, we envision that a cross-scale causal relationship from the dynamic to the cellular level serves as the basis of temperature compensability in the circadian clock system of cyanobacteria.

## Materials and Methods

### Expression and Purification of Kai Proteins

Glutathione S-transferase (GST)-tagged versions of Kai proteins were constructed in pGEX-6P-1. Each Kai GST-fusion protein was expressed in *E. coli*. BL21(DE3) and purified as reported previously ^11, 26^.

### ATPase Assay

ATPase activities of KaiC^WT^ and its mutants, dissolved in an H_2_O buffer (H1-buffer) including 20 mM Tris/HCl (pH 8.0), 150 mM NaCl, 5 mM MgCl_2_, 1 mM DTT, 1 mM EDTA, and 1 mM ATP, were measured at 303, 308, 313, and 318 K as previously reported ^11, 15^.

### *In vitro* Rhythm Assay

P-cycles of KaiC^WT^ and its mutants (0.2 mg/ml) were initiated by addition of KaiA (0.04 mg/ml) and KaiB (0.04 mg/ml) in H1-buffer at 303, 308, 313, and 318 K ^25^. For KaiC^F247A^ and KaiC^Q361E^, KaiA was added at a 2-fold higher concentration (0.08 mg/ml). Aliquots taken from the incubated samples were subjected to SDS-PAGE analysis. The relative abundances of four phosphorylation states of KaiC were quantified by densitometric image analysis of gel bands using the LOUPE software ^63^.

### Sample Preparation for QENS

Every sample was prepared on site immediately before QENS measurements. KaiC^WT^ and its mutants were purified on a gel-filtration column (Superdex 200 15/30, Cytiva) equilibrated with an H_2_O buffer (H2-buffer) containing 50 mM Tris/HCl (pH 8.0), 150 mM NaCl, 5 mM MgCl_2_, 3 mM DTT, 1 mM EDTA, and 20 mM ATP. Collected fractions were subjected to rapid buffer exchange in a D_2_O buffer (D2-buffer) containing 50 mM Tris/HCl (pD 7.6), 150 mM NaCl, 5 mM MgCl_2_, 3 mM DTT, 1 mM EDTA, and 20 mM ATP using a desalting column (HiPrep 50, Cytiva). KaiC^WT^, KaiC^S258A^, KaiC^F247A^, and KaiC^Q361E^ were concentrated up to 17.0, 12.3, 14.6, and 14.3 mg/mL, respectively. Each 1-mL sample was placed in a double-cylindrical aluminum cell with a sample thickness of 0.5 mm and sealed tightly with indium wire.

### QENS Experiments

QENS data were recorded using the near-backscattering spectrometer installed at beamline BL02 (DNA) in the Material and Life Science Experimental Facility of Japan Proton Accelerator Research Complex (MLF/J-PARC), Tokai, Ibaraki, Japan ^30^. QENS spectra were recorded over an energy transfer range from −0.5 to 1.5 meV with energy resolution (12 μeV) enabling us to access motions faster than approximately 55 ps. QENS data were collected at 302 and 312 K, or 302 and 309 K with exposure times of 6–10 h for each condition (537−615 kW). The obtained *S*(*Q,E*) were corrected for detector efficiency using a vanadium standard, and intensities were normalized as relative intensities using the standard after subtracting the contributions of the empty cell. The background spectrum of the D2-buffer was subtracted from each sample spectrum on the basis of scaling factors calculated from neutron scattering cross-sections, as reported previously ^33, 34^. The temperature was controlled by an LS350 (Lakeshore) with He conductance gas through a GM refrigerator and heat transfer from a cartridge heater installed in a copper block at the top of a sample cell, while monitoring the temperature at the bottom of the cell.

### Estimation of the Q10 values

The Q10_ATP_ and Q10_fp_ values at 303 K were estimated from the slopes of Arrhenius plots of *f*_P_ and ATPase activity, respectively, at four different temperatures (303, 305, 308, and 313 K), as previously reported ^63^. Other Q10 values were presented as the ratio of measurement results (*R*_1_ and *R*_2_) at two temperatures (*T*_1_ and *T*_2_) using 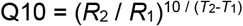.

### Calculation of *D*_global_

The *D*_global_ values of KaiC^WT^ were simulated at 302 and 312 K using a previously reported method ^42^, the crystal structure (2GBL) ^32^, and HYDROPRO ^64^ (*SI Appendix*, Section S.1.).

## Acknowledgments

We thank Dr. K. Shibata for his kind support for the trial QENS experiment, and Dr. M. Kataoka and Dr. H. Kamikubo for their discussions and critical comments on the manuscript. This study was partly supported by Grants-in-Aid for Scientific Research (17H06165 to S.A.). The QENS experiments using BL02 (DNA) at the Materials and Life Science Experimental Facility of the J-PARC were performed under user programs (Proposal No. 2017B0123, 2018B0244, 2019A0308, and 2020B0073).

