## Supplementary material for "Cross-scale Analysis of Temperature Compensation in the Cyanobacterial Circadian Clock System": SI Appendix

[a]Research Center of Integrative Molecular Systems, Institute for Molecular Science, National Institute of Natural Sciences, 444-8585 Okazaki, Japan. [b]Department of Functional Molecular Science, SOKENDAI (The Graduate University for Advanced Studies), 444-8585 Okazaki, Japan. [c]Neutron Science and Technology Center, Comprehensive Research Organization for Science and Society (CROSS), 162-1 Shirakata, Tokai, Ibaraki 319-1106, Japan. [d]Institute for Quantum Life Science, National Institutes for Quantum and Radiological Science and Technology, 2-4 Shirakata, Tokai, Ibaraki 319-1106, Japan. [e]Japan Atomic Energy Agency, 2-4 Shirakata, Tokai, Ibaraki 319-1195, Japan. [1]Contributed equally to this work. [2]Present Address: #1: Laboratoire Interdisciplinaire de Physique (LiPhy), Grenoble-Alpes University, 140 rue de la physique, 38402 Saint Martin d'Hères, France. #2: Institut Laue-Langevin, 71 avenue des Martyrs, CS 20156, 38042 Grenoble Cedex 9, France.

*Shuji Akiyama


*Satoru Fujiwara


**This PDF file includes:**

Supplementary text
Figures S1 to S4
Tables S1
SI References

### S.1 Extended Methods: Simulation of apparent diffusion coefficients (*$D_{global}$*)

$D_{global}$ of KaiC$^{WT}$ was simulated using a method originally developed by Pérez *et al* [1]. Missing regions in the crystal structure of KaiC (accession code: 2GBL) [2] were modelled using MODELLER [3]. QENS spectra, $S_{T+R}(Q,E)$, arise from contributions of both translational and rotational diffusions of the protein molecule:

$$S_{\mathrm{T+R}}(Q,E) = \sum_{l=0}^{\infty} \int_{r=0}^{D_{\mathrm{max}}} (2l+1)\rho(r) j_l^2(Qr) dr \frac{D_{\mathrm{T}}Q^2 + l(l+1)D_{\mathrm{R}}}{E^2 + (D_{\mathrm{T}}Q^2 + l(l+1)D_{\mathrm{R}})^2}$$

where $j_l(Qr)$ is the $l$th-order spherical Bessel function of the first kind; $\rho(r)$ is the radial density distribution of hydrogen atoms in the model structure; $D_{max}$ is the maximum dimension (115.3 Å) determined from the $\rho(r)$ function; and $D_T$ and $D_R$ are translational and rotational diffusion coefficients, respectively, both of which are estimated with HYDROPRO [4] using a radius of 5.6 Å for an ensemble of overlapping spheres [5]. $S_{T+R}(Q,E)$ were simulated at 302 and 312 K using $D_2O$-solvent viscosities of 1.002 and 0.806 cP, respectively [6], and then fitted using a Lorentzian function. Simulated $D_{global}$ values were determined as the linear $Q^2$-dependence of the half width at half maximum of the resultant Lorentzian function and are compiled in Table S1.

### S.2 Extended Methods: EISF analyses using diffusion-inside-two-spheres model

EISF of KaiC$^{WT}$ and the mutants were plotted against Q (Fig. S3) and then fitted using the following equation, a model containing two diffusive motions in confined spheres with distinct radii [7, 8, 9]:

$$\mathrm{EISF}(Q) = p_0 + p_1 \{ 3j_1 (Qa_1) / (Qa_1) \}^2 + p_2 \{ 3j_1 (Qa_2) / (Qa_2) \}^2 \quad [\mathbf{3}]$$

where $p_0$ is the fraction of atoms whose motions are outside the current instrumental energy window and therefore appear immobile; $p_1$ and $p_2$ correspond to the fractions of atoms diffusing within spheres of radii of $a_1$ and $a_2$, respectively; and $j_1$ denotes the spherical Bessel function of the first kind of order. Obtained parameters are summarized in Fig. S3.

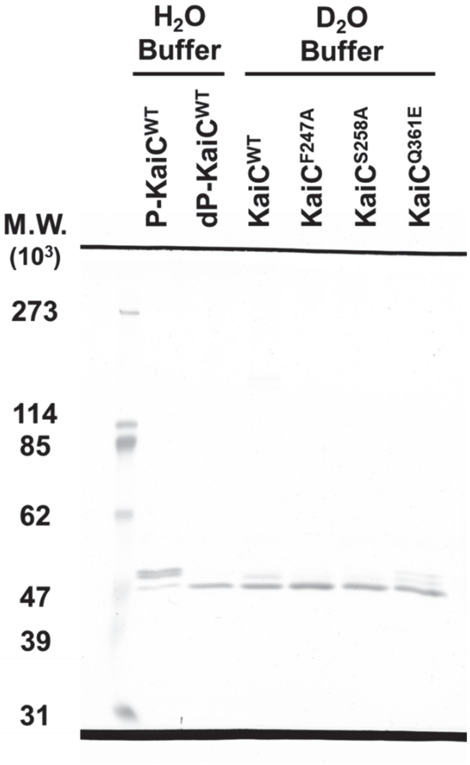


**Fig. S1.** SDS-PAGE analysis of KaiC$^{WT}$ and its mutants subjected to QENS measurements in a $D_2O$ buffer. As a reference for band assignments, phosphorylated (P-KaiC$^{WT}$) and dephosphorylated KaiC$^{WT}$ (dP-KaiC$^{WT}$) were prepared separately by incubation at 277 and 313 K, respectively, in an $H_2O$ buffer. Fractions of dP-KaiC$^{WT}$, dP-KaiC$^{F247A}$, dP-KaiC$^{S258A}$, and dP-KaiC$^{Q361E}$ were estimated by densitometric image analysis of CBB-stained gel bands to be 0.89, 0.94, 0.91, and 0.78, respectively.

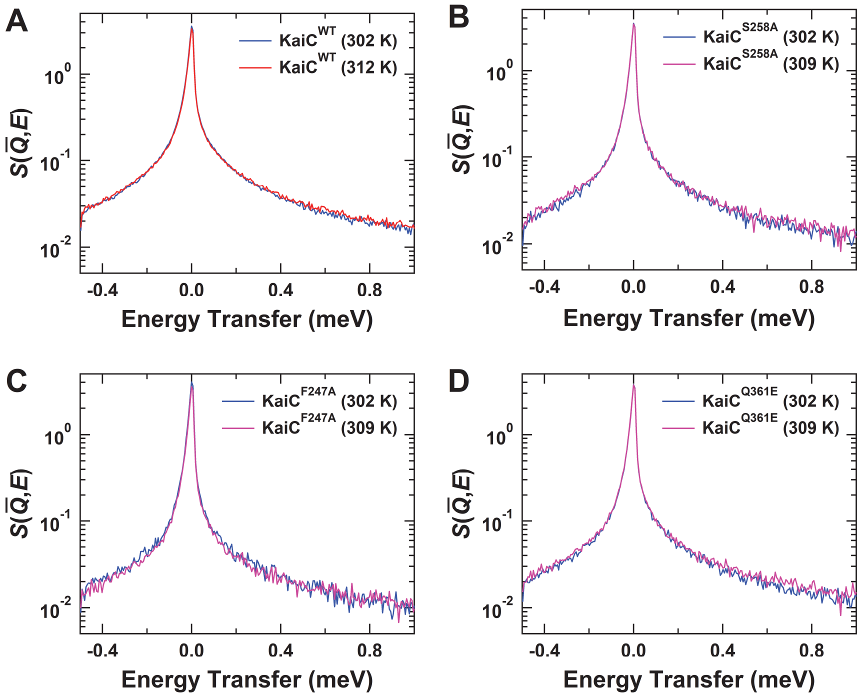


**Fig. S2.** *Q*-averaged QENS spectra, $S(\overline{Q},E)$, of (*A*) KaiC$^{WT}$, (*B*) KaiC$^{S258A}$, (*C*) KaiC$^{F247A}$, and (*D*) KaiC$^{Q361E}$.

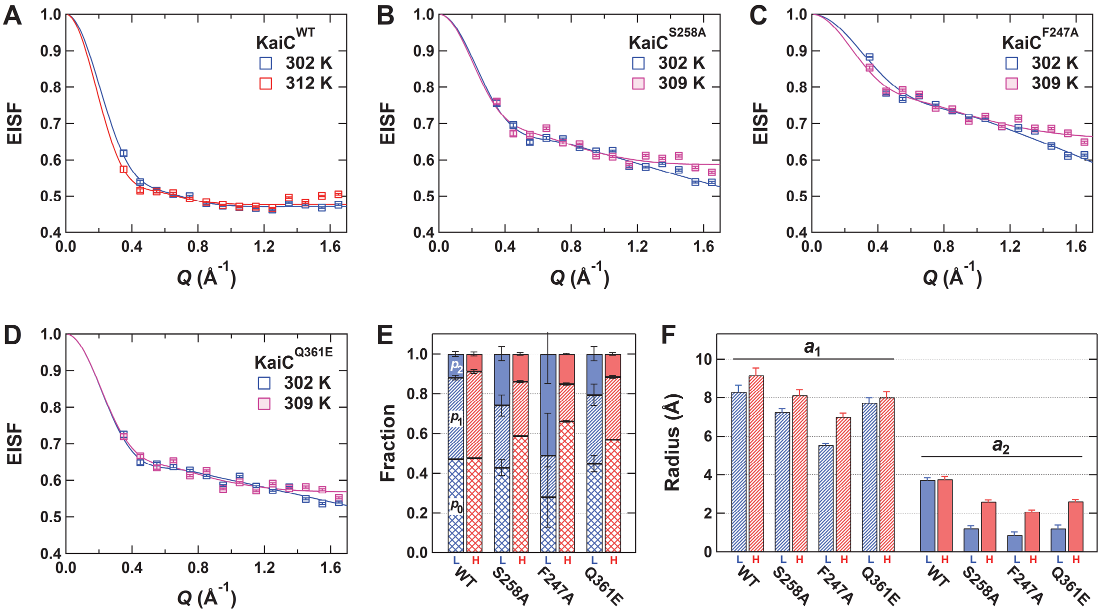


**Fig. S3.** EISF analyses under the assumption of diffusion-inside-two-spheres model for (*A*) KaiC$^{WT}$, (*B*) KaiC$^{S258A}$, (*C*) KaiC$^{F247A}$, and (*D*) KaiC$^{Q361E}$. Blue, red, and magenta boxes correspond to data acquired at 302, 312, and 309 K, respectively. Solid lines represent the resultant fits of Eq. **3**. (*E*) Immobile $p_0$ (crossed lines), mobile $p_1$ (diagonal lines), and mobile $p_2$ (filled) fractions. Blue- and red-colored bars correspond to the parameters for low (L) and high (H) temperatures, respectively. (*F*) Radii ($a_1$ and $a_2$) of spheres within which mobile atoms diffuse.

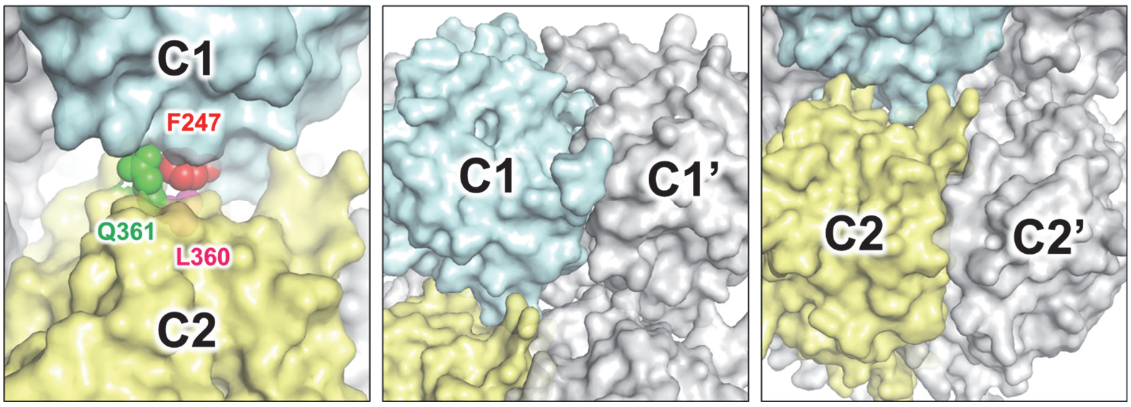


**Fig. S4.** Comparison of packing density among C1–C2 (*Left*), C1–C1' (*Center*), and C2–C2' interfaces (*Right*) in KaiC.

**Table S1.** Translational ($D_T$), rotational ($D_R$), and apparent diffusion coefficients ($D_{global}$) simulated for KaiC$^{WT}$.

| Temperature (K) | $D_T$ (cm$^2$ s$^{-1}$) | $D_R$ (s$^{-1}$) | $D_{global}$ (cm$^2$ s$^{-1}$) |
|---|---|---|---|
| 302 | 3.17 × 10$^{-7}$ | 4.89 × 10$^5$ | 3.81 × 10$^{-7}$ |
| 312 | 4.07 × 10$^{-7}$ | 6.28 × 10$^5$ | 4.88 × 10$^{-7}$ |
